# Structural basis of FANCD2 deubiquitination by USP1-UAF1

**DOI:** 10.1101/2020.12.05.412924

**Authors:** Martin L. Rennie, Connor Arkinson, Viduth K. Chaugule, Rachel Toth, Helen Walden

## Abstract

Ubiquitin-Specific Protease 1 (USP1), together with the cofactor UAF1, acts during DNA repair processes to specifically to remove mono-ubiquitin signals. The mono-ubiquitinated FANCI-FANCD2 heterodimer is one such substrate and is involved in the repair of DNA interstrand crosslinks via the Fanconi Anemia pathway. Here we determine structures of human USP1-UAF1 with and without ubiquitin, and bound to mono-ubiquitinated FANCI-FANCD2 substrate. Crystal structures of USP1-UAF1 reveal plasticity in USP1 and key differences to USP12-UAF1 and USP46-UAF1. A cryoEM reconstruction of USP1-UAF1 in complex mono-ubiquitinated FANCI-FANCD2, highlights a highly orchestrated deubiquitination process with USP1-UAF1 driving conformational changes in the substrate. An extensive interface between UAF1 and FANCI, confirmed by mutagenesis and biochemical assays, provides a molecular explanation for their requirement despite neither being directly involved in catalysis. Overall, our data provide molecular details of USP1-UAF1 regulation and substrate recognition.

## Introduction

Ubiquitination is a reversible post-translation modification involving the attachment of ubiquitin, which acts as a signal to regulate numerous cellular pathways, including protein degradation, DNA replication, and DNA repair^1,2^. The attachment of ubiquitin proceeds via a cascade of enzymes and typically involves formation of an isopeptide bond between a lysine on the substrate the carboxy-terminus of ubiquitin. Removal of the ubiquitin signal is achieved by a group of proteases known as deubiquitinases.

Ubiquitin signals are structurally diverse^1,2^, composed of either poly-ubiquitin linkages, where different ubiquitin lysines are sequentially modified, or mono-ubiquitination, in which a single ubiquitin is attached to the target protein. Ubiquitin(s) can be directly recognised by reader proteins and/or result in direct conformational changes of the modified target protein for downstream effects. Signal reversal by deubiquitinases involves recognition of these different substrate types in a spatio-temporal manner and the regulation of deubiquitinase activity^2,3^.

The largest family of deubiquitinases are the Ubiquitin-Specific Proteases (USPs). DNA repair mechanisms including the Fanconi Anemia pathway utilize USP1 for deubiquitination of mono-ubiquitinated targets^4–8^. Loss of USP1 leads to dysfunctional DNA repair and genomic instability^5,7,8^, and USP1 has been identified as a potential drug target for overcoming cancer resistance to DNA damaging drug treatment^9–11^. While we now understand how certain deubiquitinases specifically target ubiquitin-ubiquitin linkages, how USP1 targets a specific set of ubiquitin-substrate linkages remains an open question.

Three key substrates of USP1 are PCNA, FANCI, and FANCD2, each of which is mono-ubiquitinated at a specific lysine residue during repair of DNA damage^5,6,12,13^. FANCI and FANCD2 can form a heterodimer, while PCNA forms a homotrimer, with both complexes able to form DNA clamps^14–16^. FANCI and FANCD2 are homologous proteins, each with an α-solenoid fold contorted into a saxophone shape^17^. The non-ubiquitinated FANCI-FANCD2 complex is stabilized by extensive interactions across the N-Terminal Domains (NTDs) of each protein that partially bury the ubiquitination site. The C-Terminal Domains (CTDs) extend away from this interface to make an open trough, with the Head Domain connecting the NTD and CTD. Recently, cryogenic electron microscopy (cryo-EM) reconstructions of mono-ubiquitinated FANCI-FANCD2 revealed a closed conformation of the heterodimer, resulting in it encircling DNA^15,16,18^. The ubiquitin conjugated to each subunit makes non-covalent contacts with the NTD of the other subunit, reconfiguring the NTD-NTD interface and creating a CTD-CTD interface. Mono-ubiquitination of the FANCD2 subunit alone is sufficient to induce this closed conformation and results in an enhanced affinity for double stranded DNA^15,16,18,19^. The ubiquitin signal has therefore been proposed to protect DNA during repair or may recruit other repair factors. How this specific ubiquitin signal is removed to unclamp FANCI-FANCD2 post-repair is not well understood.

USP1 contains a predicted Ubiquitin-Specific Protease (USP) catalytic fold with two large insertions and an N-Terminal Extension (NTE)^20^. USP1-Associated Factor (UAF1), the catalytically essential cofactor of USP1, as well as USP12 and USP46,^21,22^ is composed of a β-propeller domain, an ancillary domain, and a SUMO-like domain (SLD). The β-propeller directly interacts with both USP12^23,24^ and USP46^25,26^ in the same way, a feature likely conserved for USP1^23^. The SLD is thought to mediate substrate recognition via binding to SUMO-Like domain Interacting Motifs (SLIMs) on substrates^23,25,27^. However, unlike USP1-UAF1, neither USP12-UAF1 nor USP46-UAF1 target mono-ubiquitinated FANCI-FANCD2^22^. Furthermore, FANCI buries a substantial surface of FANCD2’s ubiquitin^15,16^, which shields it from several deubiquitinases but not USP1-UAF1 *in vitro*^18^. Instead, FANCI phosphorylation is reported to impede USP1-UAF1 activity^28,29^; the underlying mechanism of this regulation remains unclear. Recently, biochemical work has shown that the NTE of USP1 is critical for specific removal of FANCD2’s ubiquitin^20^. However a detailed understanding of how this specialized deubiquitinase targets ubiquitin sequestered within the FANCI-FANCD2 complex is hampered by the lack of structural information.

Here we present structures of human USP1-UAF1 alone, bound to ubiquitin, and finally bound to a mono-ubiquitinated FANCI-FANCD2 substrate, representing the first human deubiquitinase-substrate complex, providing insight into the catalytic cycle of USP1-UAF1. Crystal structures of truncated USP1-UAF1, with and without ubiquitin, reveal features shared with USP12/46-UAF1 complexes, and those that aid USP1’s isopeptidase activity. A cryo-EM structure of full-length USP1-UAF1 bound to FANCI and mono-ubiquitinated FANCD2 reveals an extensive UAF1-FANCI interface that surprisingly does not involve the previously predicted FANCI SLIM. Conformational changes are observed in each entity of the deubiquitinase-substrate complex, in particular FANCD2’s ubiquitin disengages from FANCI and associates with USP1. In addition, the structures reveal how engagement of USP1-UAF1 can occur while the FANCI-FANCD2 heterodimer is clamped on DNA. Using mutational analyses and biochemical assays we highlight the importance of the interfaces within the structures. Overall the structures and assays clarify a highly orchestrated and multifaceted deubiquitination cycle, in which two binding partner proteins, UAF1 and FANCI, along with USP1 contacts with FANCD2, regulate the reversal of this critical ubiquitin signal.

## Results

### Architecture and plasticity of USP1

In order to understand how USP1 is regulated by UAF1 and how USP1 targets ubiquitin, we sought to determine structures of the multiple stages in USP1’s catalytic cycle. We determined structures of USP1-UAF1 with and without ubiquitin using X-ray crystallography. Full-length constructs of USP1-UAF1 did not yield well-diffracting crystals. In order to obtain USP1-UAF1 crystals that diffract to better than 4 Å, we deleted several flexible regions of USP1, predicted to be disordered, and the SLD of UAF1 (Fig. 1a, Methods). We determined a ubiquitin-bound structure of the same construct by reacting USP1 with propargylated ubiquitin^30^ (Fig. 1b). The USP1 structure exhibits a typical “fingers-palm-thumb” architecture (Fig. 1a). Ubiquitin sits in the palm, making contacts with all three subdomains, while only the fingers interact with UAF1’s β-propeller domain, as observed for USP12-UAF1^23,24^ and USP46-UAF1^25^. Small angle X-ray scattering (SAXS) measurements confirmed the oligomeric arrangement observed in the crystal structures (Fig 1c, Supplementary Fig. S1a-c). The deleted Inserts 1 and 2 are positioned on the opposite face of USP1 to the ubiquitin binding site, while the deleted N-terminus extends from the distal tip of USP1, furthest from UAF1 (Fig. 1B).

**Fig. 1.**
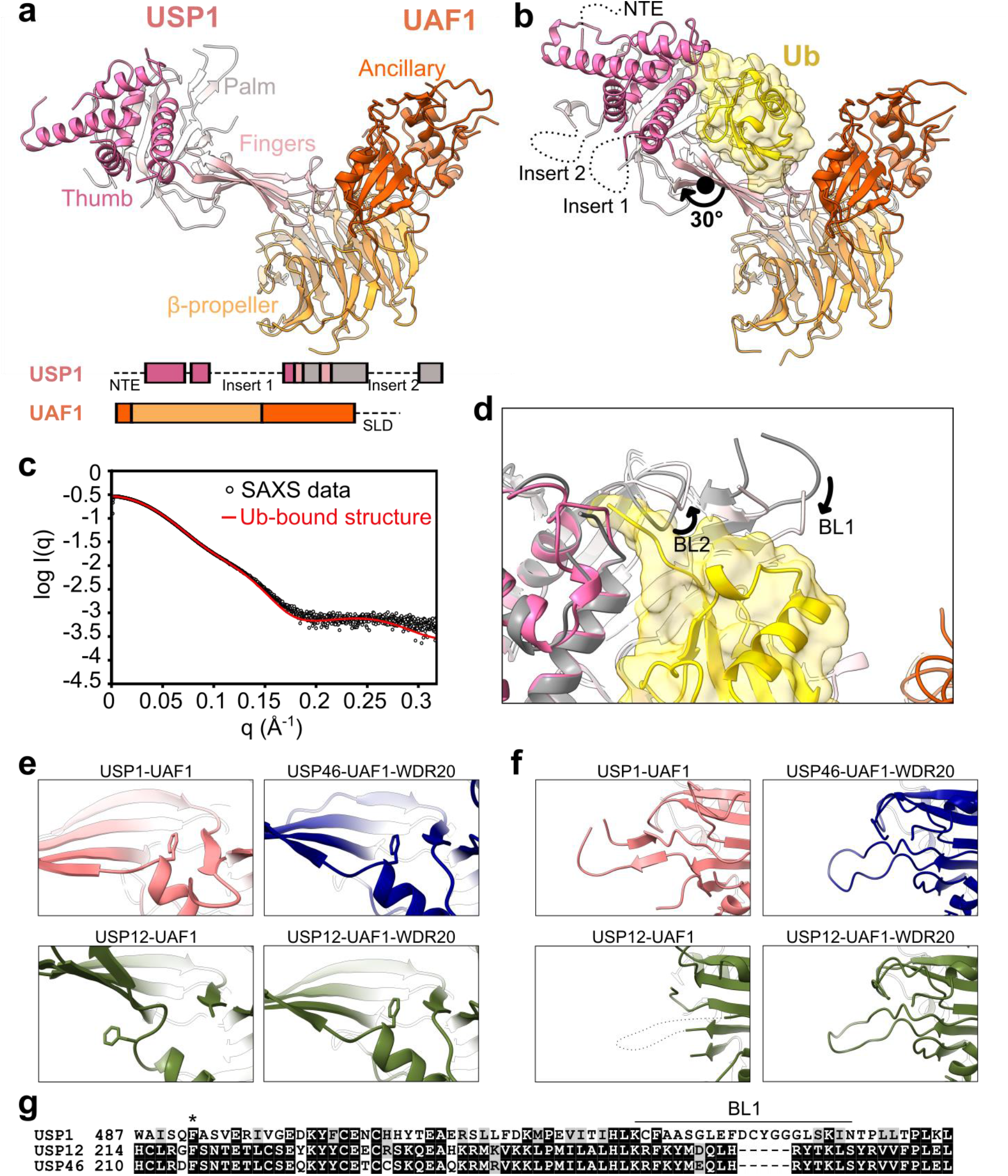
Structural characterization of USP1-UAF1. (**a**) Crystal structure of USP1, lacking the NTE and Insert 1 and Insert 2, in complex with UAF1 lacking the SLD. Subdomains of USP1 and domains of UAF1 are indicated. Deleted regions are indicated in the primary structure as dashed lines (see also Methods). (**b**) Crystal structure of USP1 lacking the N-terminal region and Insert 1 and Insert 2 covalently modified with propargylated ubiquitin in complex with UAF1 lacking the SLD. The termini and approximate positions where the NTE, Insert 1, and Insert 2 would be are indicated. The rotation axis relating the palm and thumbs of the two structures is shown. (**c**) Fit of the ubiquitin-bound crystal structure with solution state SAXS measurements. (**d**) Comparison of USP1 catalytic region between the ubiquitin-free (gray) and ubiquitin-bound (colored) crystal structures. Structures were aligned by the USP1 subunit. (**e**) A phenylalanine, conserved in USP1, USP12, and USP46, occupies a pocket between the palm and fingers in ubiquitin-free USP1-UAF1 structure, and in USP46-UAF1-WDR20 (6JLQ)^26^ and USP12-UAF1-WDR20 (5K1C) but not USP12-UAF1 (5K1A)^23^. (**f**) The BL1 is ordered in ubiquitin-free USP1-UAF1 structure, and in USP46-UAF1-WDR20 (6JLQ)^26^ and USP12-UAF1-WDR20 (5K1C) but not USP12-UAF1 (5K1A)^23^. (**g**) Multiple sequence alignment of USP1, USP12, and USP46 in the region including the conserved phenylalanine (indicated by *) and the BL1. Sequence alignment was performed using Clustal Omega (http://www.ebi.ac.uk/Tools/msa/clustalo/) and visualized using BOXSHADE.

Comparison of the ubiquitin bound and unbound structures of USP1-UAF1 reveals conformational changes in USP1. In the ubiquitin-bound structure, “Binding Loops” BL1 and BL2 are re-arranged with respect to the ubiquitin-free structure, which allows threading of the ubiquitin tail between the palm and thumb (Fig. 1d), while the globular body of ubiquitin primarily contacts the fingers, similar to other USPs. In addition to these local changes, USP1 exhibits flexibility in the fingers, with a 30° rotation relating the palm and thumb of the two crystal structures (Fig. 1a-b). Furthermore, each crystal had two USP1-UAF1 complexes per asymmetric unit both with similar conformations within the same asymmetric unit (Supplementary Fig. S1d). Such structural plasticity has been also observed for crystal structures of USP12-UAF1 and USP46-UAF1 and may facilitate substrate engagement.

Although USP1, USP12, and USP46 are all regulated by UAF1, only USP12 and USP46 activities are enhanced by another cofactor, WDR20^31,32^. Several structural features of the ubiquitin-free USP1-UAF1 structure correspond more closely to WDR20-bound USP12 and USP46 structures than WDR20-free USP12^23,26^ (Fig. 1e-g). A conserved phenylalanine is buried between the fingers and palm, for USP1-UAF1, USP12/46-UAF1-WDR20, but not USP12-UAF1 (Fig. 1e). Correlating with this, USP1’s BL1 is ordered in USP1-UAF1, USP12/46-UAF1-WDR20 but not USP12-UAF1 (Fig. 1f), however crystal packing effects of the ubiquitin-free USP1-UAF1 structure may contribute to this stabilization. Therefore, the USP fold of USP1 may be primed for efficient catalysis without the additional requirement of WDR20. Overall, our crystal structures reveal typical USP-ubiquitin and USP-UAF1 interactions and highlight both the differences between USP1 and USP12/46 along with the positioning of the inserts and NTE of USP1.

### Structure of USP1-UAF1 bound to FANCI-FANCD2 substrate

Our USP1-UAF1 structures allow comparisons to USP12/46 complexes, demonstrate plasticity in the USP fold and show of USP1 binds ubiquitin. However, in order to understand how USP1-UAF1 recognises mono-ubiquitinated substrate, we determined a cryo-EM structure of full length USP1-UAF1 bound to the FANCI-FANCD2^Ub^-DNA substrate with ubiquitin only on FANCD2. This singly mono-ubiquitinated substrate is readily deubiquitinated by USP1-UAF1 even in the presence of DNA, unlike the doubly mono-ubiquitinated complex, FANCI^Ub^-FANCD2^Ub 18,20,33^. We employed a catalytically compromised C90S mutant of USP1, and our previous strategy for generating the mono-ubiquitinated substrate^34,35^ to reconstitute the USP1^C90S^-UAF1-FANCI-FANCD2^Ub^ complex, with full length versions of all four proteins (Supplementary Fig. S2a). We then determined a cryo-EM reconstruction of this enzyme-substrate complex, bound to double-stranded DNA, at 3.8 Å global resolution (FSC=0.143) and local resolution 3.6-10 Å (FSC=0.5) (Fig. 2a and Supplementary Fig. S2b-d). Using our crystal structure of ubiquitin-bound USP1-UAF1 described above (Fig. 1), and the structures of mono-ubiquitinated human FANCI-FANCD2^Ub^ (PDB ID: 6VAF) and full-length UAF1 (PDB ID: 5K1A) we built a model of approximately 70% of the residues in the complex including the K561-G76 isopeptide linkage (Fig. 2b-c).

**Fig. 2.**
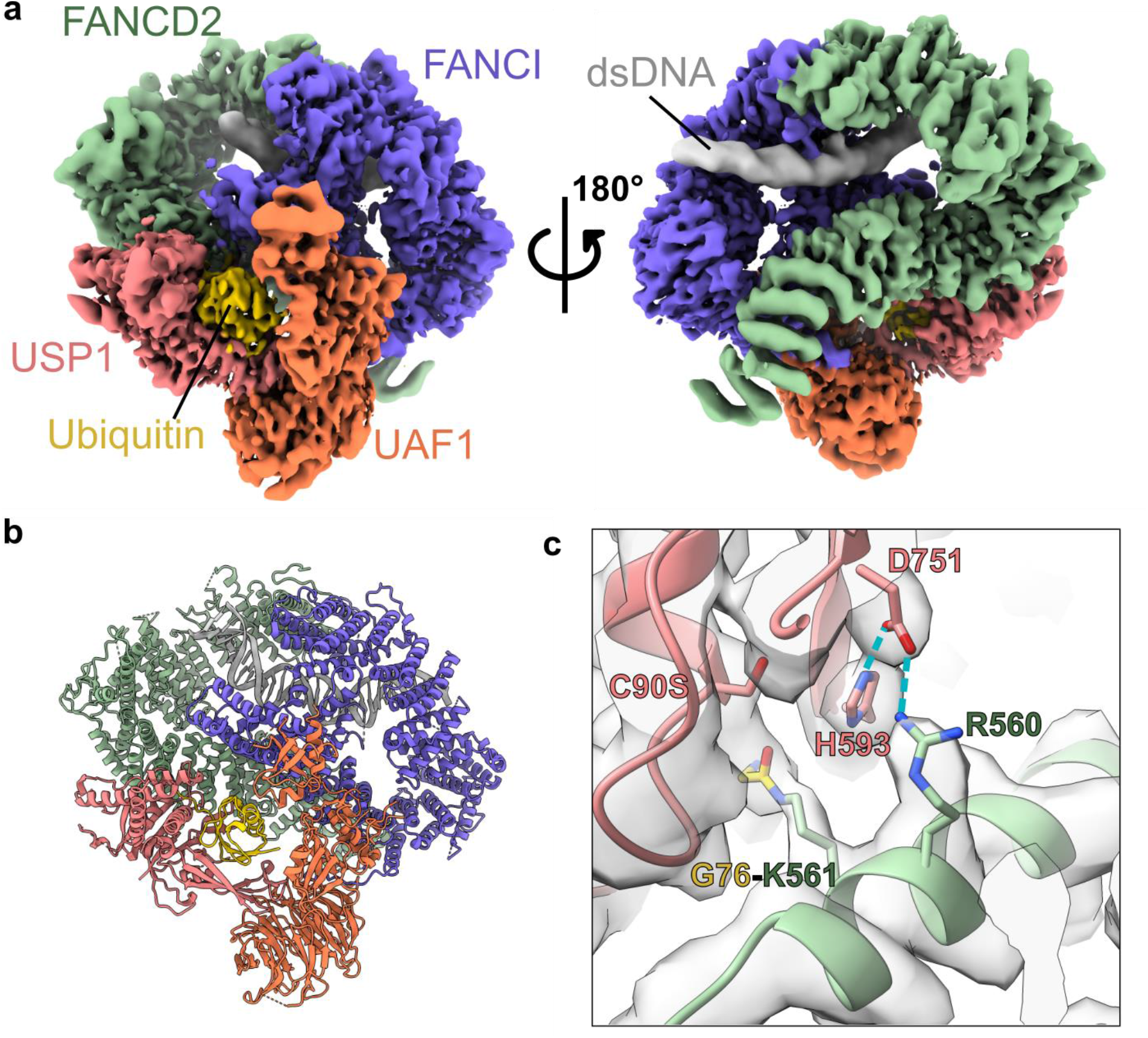
Structural characterization of full-length USP1-UAF1 bound to the FANCI-FANCD2^Ub^ substrate. (**a**) Locally filtered cryo-EM map of the complex containing USP1 (pink), UAF1 (orange), FANCI (violet), and ubiquitinated (yellow) FANCD2 (green) at a threshold of 0.07. (**b**) Model of the enzyme-substrate complex. (**c**) The catalytic site of USP1 with the FANCD2^K561^-ubiquitin^G76^ isopeptide linkage overlaid with the DeepEMhancer^36^ map at a threshold of 0.1. Hydrogen bonds are shown as blue dashed lines.

Globally, USP1-UAF1 and the ubiquitin of FANCD2 are in a similar arrangement to the ubiquitin-bound crystal structure described above (Fig. 1). FANCI-FANCD2 are in a closed conformation encircling DNA, similar to that observed for the singly and doubly monoubiquitinated complexes^15,16,18^. The enzyme-substrate complex is arranged such that USP1 contacts FANCD2 while UAF1 contacts FANCI in a 1:1:1:1 stoichiometry.

### UAF1 and FANCI form a scaffold for deubiquitination

UAF1 and FANCI have an extensive interface that involves interactions from the β-propeller, ancillary, and SLD of UAF1 and the FANCI NTD and HD, and buries ∼1600 Å^2^ (Fig. 3a). Interestingly, a FANCI SLIM^27^, previously proposed to mediate interactions with UAF1, was not located at the interface observed here. Instead, a FANCI loop (^251^DELLDVV^257^) directly interacts with the SLD of UAF1 (Fig. 3a), with L254 inserting into the SLD hydrophobic groove. The UAF1-FANCI interface observed here likely explains how FANCI favours USP1-UAF1 targeting over more promiscuous deubiquitinases^18^.

**Fig. 3.**
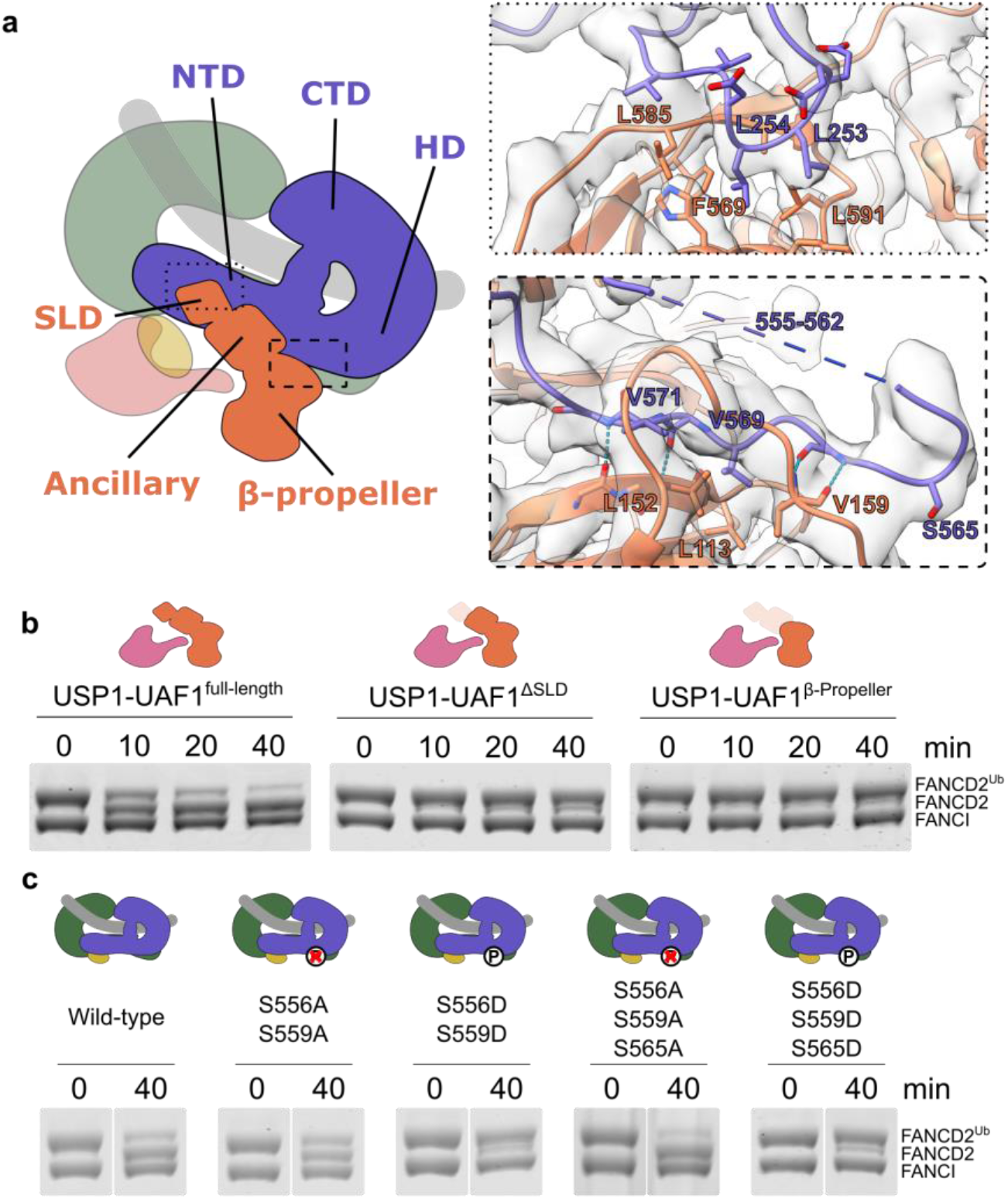
FANCI-UAF1 interactions are important for deubiquitination of FANCI-FANCD2^Ub^. (**a**) The interface between FANCI (violet) and UAF1 (orange). The FANCI^249-260^ loop inserts into the hydrophobic pocket of the SLD of UAF1, while the FANCI^547-576^ region interacts with the β-propeller domain of UAF1. Hydrophobic side chains involved in the interaction are shown and backbone hydrogen bonds are highlight as blue dashed lines. (**b**) Truncations of UAF1 reduce FANCI-FANCD2^Ub^ deubiquitination. Deubiquitination of FANCD2^Ub^ (1 μM) in complex with non-ubiquitinated FANCI (1 μM) by full-length USP1 reconstituted with UAF1 or truncations of UAF1 (200 nM enzyme complex) was assessed by SDS-PAGE and Coomassie staining (see also Supplementary Fig. S3). (**c**) Phosphorylation within FANCI^547-576^ regulates deubiquitination. Deubiquitination of FANCD2^Ub^ (1 μM) in complex with non-ubiquitinated FANCI phosphorylation mimics (1 μM) by full-length USP1 reconstituted with full-length UAF1 (100 nM enzyme complex) was assessed using SDS-PAGE and Coomassie staining. All assays were in the presence of 4 μM 61 base pair dsDNA, and performed at least twice (two technical replicates).

To further test this hypothesis, we compared the deubiquitination activity of USP1 reconstituted with either full-length UAF1, or UAF1 truncations lacking the SLD (UAF1^ΔSLD^) or comprising just the β-propeller domain (UAF1^β-propeller^). Both truncations severely impaired deubiquitination of FANCI-FANCD2^Ub^ compared to full-length UAF1 (Fig. 3b). To confirm that this was not due generic reduction in activity we also assessed deubiquitination of FANCD2^Ub^ in the absence of FANCI (Supplementary Fig. S3a-d). For the UAF1 truncations deubiquitination of FANCD2^Ub^ alone was only slightly reduced compared to full-length, as also reported for USP46-UAF1^β-propeller^ acting on fluorescently tagged ubiquitin^25^. Indeed, even at enzyme concentrations resulting in comparable deubiquitination of FANCD2^Ub^ alone, activity against FANCI-FANCD2^Ub^ for the UAF1^β-propeller^ was reduced compared to full-length UAF1, confirming the importance of the UAF1-FANCI interaction (Supplementary Fig. 3d). In the absence of UAF1 (USP1 alone) we did not detect deubiquitination of either substrate, consistent with the requirement of UAF1 for deubiquitinase activity of USP1 (Supplementary Fig. S3a-b). Overall, these data suggest that the SLD and Ancillary domain are important for deubiquitination in the context of the FANCI-FANCD2 heterodimer.

In addition, the β-propeller domain of UAF1 interacts with a loop connecting the NTD and HD of FANCI (FANCI^547-576^), the phosphorylation of which inhibits deubiquitination by USP1-UAF1^28,29^ (Fig. 3a). Although density for the FANCI phosphorylation targets (S556, S559, and S565) is weak in our reconstruction, neighbouring residues contribute hydrophobic interactions and backbone hydrogen bonds to the interaction. We confirmed a role of FANCI phosphorylation in regulating FANCI-FANCD2^Ub^ deubiquitination using different FANCI mutants (Fig. 3c, Supplementary Fig. 3e). The addition of FANCI phosphomimic S556D-S559D or S556D-S559D-S565D mutants resulted in reduced FANCD2 deubiquitination compared to addition of wild-type FANCI. On the other hand, phosphodead S556A-S559A or S556A-S559A-S565A mutants were similar to wild-type. Finally, we note that mutation of the previously proposed SLIM motif of FANCI^27^ (I683A-L685A) resulted in similar deubiquitination to wild-type FANCI, confirming its redundancy for FANCI-FANCD2^Ub^ deubiquitination *in vitro* (Supplementary Fig. S3e). We propose that phosphorylation alters the conformation of FANCI^547-576^ such that it has reduced competency for binding to UAF1. Taken together, the extensive UAF1-FANCI interface, along with FANCI phosphorylation influence efficient USP1-UAF1 activity and reveal the multi-faceted regulation underlying FANCD2 deubiquitination.

### Ubiquitin is poised for removal from FANCD2

The ubiquitin conjugated to FANCD2 undergoes a large rearrangement in the enzyme-substrate, USP1-UAF1-FANCI-FANCD2^Ub^, complex relative to the substrate only, FANCI-FANCD2^Ub^, complex (Fig. 4a, Supplementary Movie S1). In the FANCI-FANCD2^Ub^ complex, the ubiquitin tail is bent in a conformation incompatible with USP1 binding, with ubiquitin R72 forming intramolecular interactions with FANCD2 and the ubiquitin hydrophobic patch forming intermolecular interactions with FANCI^15,16^. In the USP1-UAF1-FANCI-FANCD2^Ub^ complex, ubiquitin is rotated by 160° and extracted from its non-covalent interaction with FANCI to sit in the hand of USP1, with the ubiquitin tail extended to sit between the palm and thumb (Fig. 4a, Supplementary Fig. S4a). The resulting USP1-ubiquitin interaction is analogous to the ubiquitin-bound crystal structure above (Fig. 1b), however the palm and thumb are slightly more contracted with respect to the fingers (Supplementary Fig. S4b).

**Fig. 4.**
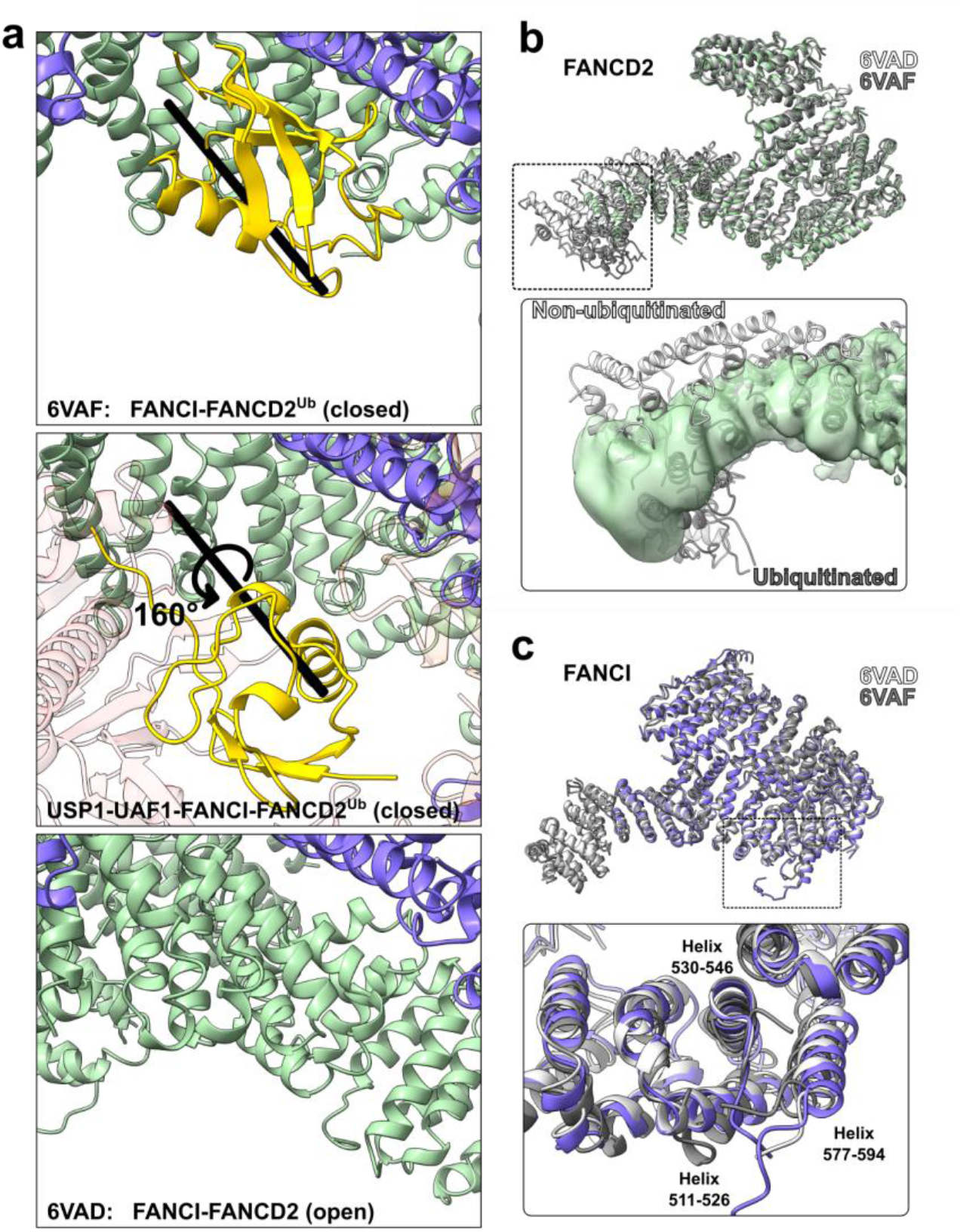
Conformational changes in FANCI (green) and FANCD2 (violet) during the deubiquitination cycle. (**a**) Conformational changes of ubiquitin (yellow) associated with USP1-UAF1 binding. Structures of the FANCI-FANCD2^Ub^ substrate of deubiquitination (6VAE) and the FANCI-FANCD2 product (6VAD)^15^ were aligned to the USP1-UAF1-FANCI-FANCD2^Ub^ structure by the FANCI subunit. The axis of rotation for ubiquitin is shown as a black cylinder. In the deubiquitinated state (bottom panel) FANCD2 and USP1 would sterically clash. (**b**) Aligned FANCD2 subunits from FANCI-FANCD2^Ub^ (6VAF; dark gray), USP1-UAF1-FANCI-FANCD2^Ub^ (green), and FANCI-FANCD2 (6VAD; light gray). The N-terminus of FANCD2 in the USP1-UAF1-FANCI-FANCD2^Ub^ structure (locally filter map at a threshold of 0.033) is in an intermediate conformation between the FANCI-FANCD2^Ub^ and FANCI-FANCD2 states. (**c**) Aligned FANCI subunits from FANCI-FANCD2^Ub^ (dark gray), USP1-UAF1-FANCI-FANCD2^Ub^ (violet), and FANCI-FANCD2 (light gray). Although all three states are globally very similar, FANCI of USP1-UAF1-FANCI-FANCD2^Ub^ more closely resembles the non-ubiquitinated FANCI-FANCD2 state in the HD close to the UAF1 interface.

While the USP1-ubiquitin interaction buries ∼1800 Å^2^, the observed USP1-FANCD2 interaction is significantly smaller, burying ∼600 Å^2^. Within the USP1-FANCD2 interface, a conserved arginine on FANCD2 (R560), adjacent to the ubiquitinated lysine, hydrogen bonds with D751 of the USP1 catalytic triad (Fig. 2c). The catalytic cysteine, mutated to serine in our structure, appears to face away from the isopeptide bond, indicating local changes in the active site are required before hydrolysis can occur.

Sitting at the crux between FANCI, FANCD2, and the BL1 of USP1 is weak density (Supplementary Fig. S4c). Given its location it could belong to the NTE of USP1, the BL1 of USP1 or the N-terminus of UAF1, however we were not able to unambiguously assign this density. Contributing to this uncertainty, the NTE density terminates adjacent to the USP domain and FANCD2 (Supplementary Fig. S4d), leaving 74 residues unmodelled. Density for Inserts 1 and 2 of USP1 is absent in our structure (Supplementary Fig. S4e), suggesting that these are not tightly associated to the USP domain and consistent with their disposability for FANCI and FANCD2 deubiquitination^20^.

### The closed conformation of FANCI-FANCD2 is recognised by USP1-UAF1

The FANCI-FANCD2 heterodimer remains in the closed conformation that encircles DNA when USP1-UAF1 is bound, despite disruption of the non-covalent interaction between FANCD2’s ubiquitin and FANCI (Fig. 2a). The open, trough shaped FANCI-FANCD2 conformation would be incompatible with the mode of USP1-UAF1 binding identified here due to steric clashes with USP1 (Fig. 4a, bottom panel). However, the FANCD2 subunit adopts an intermediate conformation to the ubiquitinated and non-ubiquitinated extremes (Fig. 4b). The NTD of FANCD2 was previously shown to flex with respect to the CTD, adopting an extended conformation in unmodified FANCD2 and bent when it is ubiquitinated^15^. When bound to USP1-UAF1, the NTD of FANCD2 becomes more extended than the ubiquitinated extreme, however retains some flexibility as illustrated by 3D variability analysis^37^ (Supplementary Movies S2-S3).

Although residues 1-180 of FANCI had weak density and were not modelled for our structure, the rest of FANCI remains in a globally similar conformation regardless of ubiquitination state or binding of USP1-UAF1 (Fig. 4c). However, there are slight changes in the region 437-790, encompassing part of the NTD and most of the HD, including the FANCI ubiquitination site. The USP1-UAF1 bound state more closely aligns with the non-ubiquitinated state, particularly for helices 511-526, 530-546, and 577-594 (Fig. 4c), which are adjacent UAF1 β-propeller binding site. Interestingly, the other face of this region contacts the FANCD2 NTD. Therefore UAF1 interaction with FANCI may allosterically regulate the interaction between FANCI and FANCD2.

## Discussion

The structures described here provide insight into the catalytic cycle of USP1-UAF1 deubiquitination and mechanisms by which the cycle is controlled (Fig. 5). The ubiquitin-free crystal structure (Fig. 1a) represents the free enzyme (E), while the cryo-EM structure (Fig. 2b) represents a substrate-bound state (ES), in which the enzyme has globally distorted the substrate. Local changes in the active site are further required to distort the isopeptide bond into the transition state before hydrolysis occurs. Subsequent dissociation of deubiquitinated FANCI-FANCD2 in the open conformation (P) likely occurs prior to dissociation of ubiquitin, as occurs for other USPs^38^. We reason that this would occur due to the narrow size of the pocket through which ubiquitin could leave the in the USP1-UAF1-FANCI-FANCD2^Ub^ structure, and the conformational changes occurring that appear to be restoring the open conformations of the FANCI and FANCD2 subunits. Finally, ubiquitin-bound enzyme (EP), represented here by the ubiquitin-bound crystal structure (Fig. 1b), must release the ubiquitin to complete catalytic cycle.

**Fig. 5.**
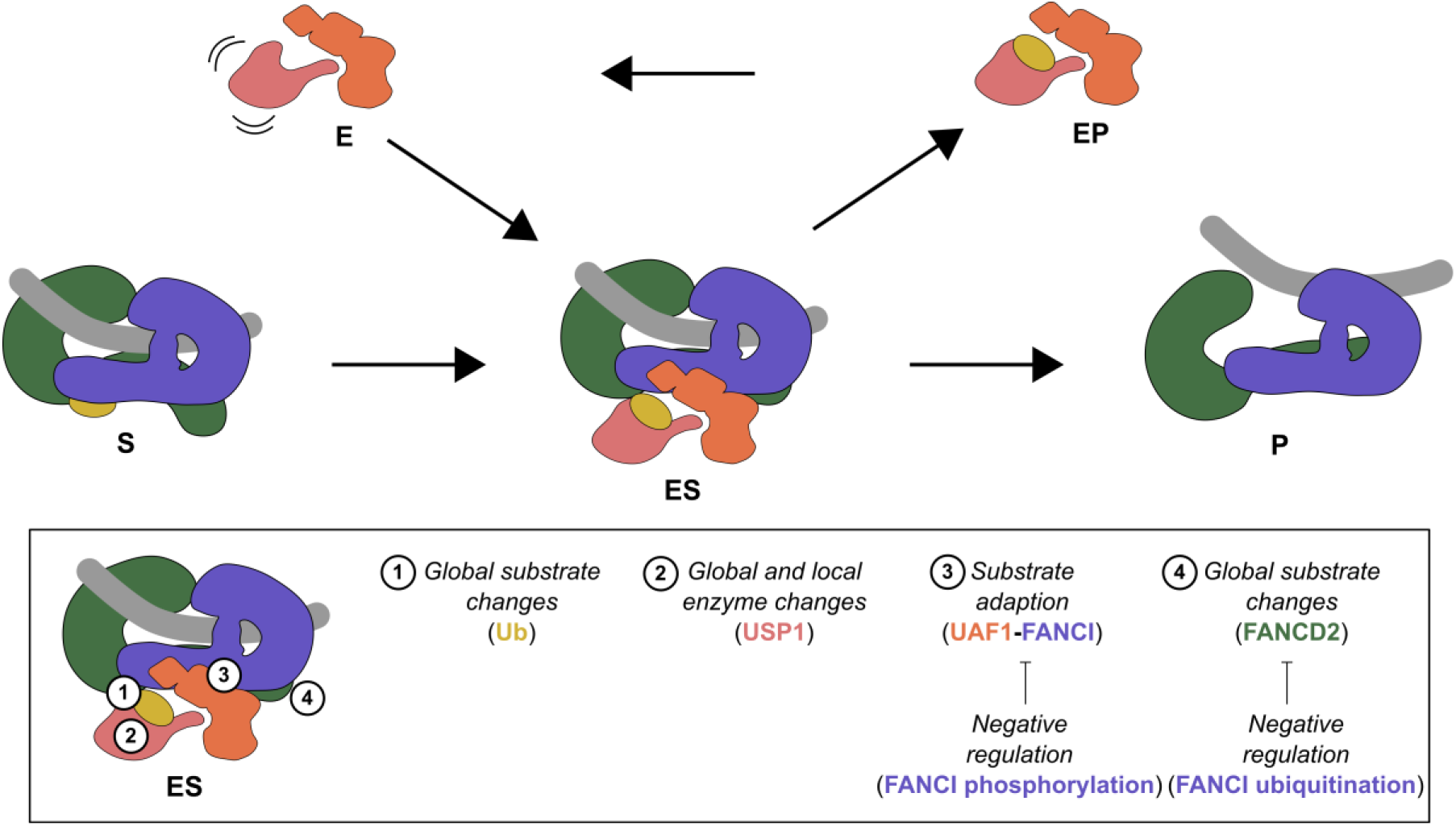
Schematic representation of the deubiquitination cycle of FANCI-FANCD2 by USP1-UAF1. Key features of the reaction are highlighted.

Enzymes acting on large proteins typically utilize various regions of the substrate and the enzyme distal to the active site, to facilitate catalysis^39^. The USP1-UAF1 enzyme and FANCI-FANCD2^Ub^ substrate are prime examples of this complexity, with regulation occurring at multiple places (Fig. 5). Furthermore, in this system, two relatively rigid subunits, UAF1 of the enzyme complex and FANCI of the substrate complex, appear to act as a scaffold, while the catalytic subunit, USP1, and the ubiquitin and conjugated protein, FANCD2, are much more flexible. Such flexibility may be an essential feature of the machinery, allowing for gross conformational changes to occur, much in the same way local rearrangements of a catalytic site are required in enzymes acting on small molecules.

Although we were unable to unambiguously locate the NTE of USP1, which we have previously shown is necessary for efficient deubiquitination of FANCD2^20^, we did identify a candidate region (Supplementary Fig. S4c). Curiously, the region identified sits between FANCI and FANCD2 with two aromatic residues in close proximity (FANCI^F277^ and FANCD2^Y520^). Both of these residues are buried by the NTD-NTD interface in the open, non-ubiquitinated FANCI-FANCD2 state. As such the occupant of the density may maintain the closed conformation while FANCI and FANCD2 return to their respective open conformations. The proximity of the BL1 of USP1, which contains a small insertion compared to USP12 and USP46 (Fig. 3g) suggests the possibility of a *cis* interaction between the NTE of USP1 and its BL1. However, further work is required to confirm these hypotheses. Unlike the NTE of USP1, which facilitates deubiquitination of FANCD2 in general, the UAF1-FANCI interaction facilitates deubiquitination in the context of the FANCI-FANCD2 heterodimer.

Our structure demonstrates that USP1 can recognize the FANCI-FANCD2 complex assembled on DNA and in the closed conformation. However, the role of DNA in regulation of deubiquitination is controversial with studies reporting DNA-mediated inhibition^18,20,33^ and enhancement^40,41^ of deubiquitination. Such discrepancies may originate from binding to FANCI-FANCD2 and USP1-UAF1, respectively. Despite addition of dsDNA after formation of the enzyme-substrate complex we did not find evidence of DNA binding to USP1-UAF1 suggesting dsDNA preferentially binds to FANCI-FANCD2. Upon dissociation of USP1-UAF1, return of the FANCI-FANCD2 to the open conformation may unlock it from DNA.

Within the USP1-UAF1-FANCI-FANCD2^Ub^ structure there is space to accommodate ubiquitin conjugated to FANCI. As such we expect a similar mechanism for recognition of the FANCI^Ub^-FANCD2^Ub^, for FANCD2 deubiquitination, as FANCI-FANCD2^Ub^. FANCD2 is deubiquitinated slower in the FANCI^Ub^-FANCD2^Ub^ complex with I44 of FANCI’s ubiquitin playing an important role^18,20,33^. The interaction between FANCI’s ubiquitin and FANCD2’s NTD^15^ may restrict flexing of the NTD observed in the USP1-UAF1-FANCI-FANCD2^Ub^ structure (Fig. 4b). Alternatively, the conjugation of ubiquitin to FANCI may reduce the propensity for USP1-UAF1 to return of the FANCI subunit to its open conformational state (Fig. 4c). Either or both of these would provide a mechanism for the protection of FANCD2 from deubiquitination that is achieved by FANCI ubiquitination. Although the order of deubiquitination of FANCI^Ub^-FANCD2^Ub^ is not known, the FANCD2 subunit is more rapidly deubiquitinated in biochemical assays and therefore expected to be deubiquitinated first. How FANCI is deubiquitinated remains an open question and will require further enzyme-substrate structures.

Although USP1 and FANCI compete for the hydrophobic patch of ubiquitin, the surface area of ubiquitin buried by FANCI (1450 Å^2^)^15^ is less than that of USP1. As the FANCI binding surface is remains accessible in our structure, the larger surface buried by USP1 may allow for preferential binding of ubiquitin to USP1 over FANCI. In the doubly mono-ubiquitinated complex, the surface area of the ubiquitin conjugated to FANCI, buried by FANCD2 is 1820 Å^2 15^. Therefore, it may be more difficult for USP1-UAF1 to extract the ubiquitin conjugated to FANCI from the complex, consistent with the observation that deubiquitination of FANCD2’s ubiquitin occurs more rapidly than FANCI’s ubiquitin^18,20,33^.

Finally, we have revealed the mechanism FANCI recognition by UAF1 (Fig. 3). Unlike the NTE of USP1, which facilitates deubiquitination of FANCD2 in general^20^, the UAF1-FANCI interface facilitates deubiquitination in the context of the FANCI-FANCD2 heterodimer. This interface appears to act as a switchable substrate adaptor, allowing USP1 to target ubiquitinated FANCD2 when FANCI is dephosphorylated. In addition it contributes to the ability of USP1 to overcome the masking of the hydrophobic patch of ubiquitin by FANCI. The FANCI region containing the S556, S559, and S565 phosphorylation sites appears to be a key player in this interaction. Interestingly, crystal structures of mouse FANCI-FANCD2 (3S4W) and FANCI alone (3S51) show this region two different conformations^17^, while in cryo-EM structures of human FANCI-FANCD2 without UAF1 the region is not well ordered (6VAA, 6VAD, 6VAE, 6VAF)^15^. Therefore, the dynamics of this region may be important for regulating recognition by UAF1. Overall, our results reveal a sophisticated deubiquitination process in which multiple regions of the substrate are recognized to coordinate both ubiquitin removal and substrate conformation.

## Methods

### Protein expression and purification

Proteins were prepared described previously^18,20,34^. Briefly, His-TEV-USP1^Δ1-66,Δ125-138,Δ229-408,Δ608-737^-UAF1^Δ564-677^ (USP1^ΔNΔ1Δ2^-UAF1^ΔSLD^) was co-expressed in *Sf*21 insect cells and purified by Ni-NTA affinity, anion exchange, TEV protease treatment, subtractive Ni-NTA, and gel filtration^18^. His_6_-TEV-USP1^G670A,G671A^, His_6_-TEV-USP1^G670A,G671A,C90S^, and His_6_-TEV-UAF1^27-359^ (β-propeller domain) were expressed in *Sf*21 cells and purified by Ni-NTA affinity, anion exchange, TEV protease treatment, subtractive Ni-NTA, and gel filtration^20^. His_6_-3C-UAF1 and His_6_-TEV-UAF1^2-580^ (ΔSLD) were expressed in *Sf*21 cells and purified by Ni-NTA affinity, anion exchange, and gel filtration (without removal of the tag)^20^. Propargylated ubiquitin was prepared from a ubiquitin-intein-CBD construct via reaction with MESNa then proparglamine at 4°C, and reacted with USP1^ΔNΔ1Δ2^-UAF1^ΔSLD^ as described previously^20^.

His_6_-3C-FANCD2, His_6_-FANCI, His_6_-TEV-V5-FANCI, and His_6_-TEV-V5-FANCI mutants were expressed in *Sf*21 cells and purified by Ni-NTA affinity, anion exchange, and gel filtration^34^. FANCD2 was ubiquitinated and purified using an engineered Ube2T and SpyCatcher-SpyTag setup^34^. The His_6_-3C tag was removed during preparation of ubiquitinated FANCD2, while for FANCI tags were not removed.

Protein concentrations were determined using the predicted extinction coefficients at 280 nm^42^ and absorbance via a Nanodrop. The ratio of 260nm/280nm was ≤0.65 for all protein batches used in subsequent experiments.

### Crystallization, data collection, and processing

Crystals used to solve the structures were grown by hanging drop at 19°C. Purified USP1^ΔNΔ1Δ2^-UAF1^ΔSLD^ was concentrated to 11-15 mg/mL and drops of 4.5 μL were setup using a reservoir of 10% w/v PEG4000, 100 mM MES/Imidazole pH 6.0-6.5, 30 mM CaCl_2_, 30 mM MgCl_2_, 20-25% glycerol v/v and a ratio of 2 volumes protein to 1 volume reservoir. Purified USP1^ΔNΔL1ΔL2^-UAF1^ΔSLD^ was reacted with propargylated ubiquitin (USP1^ΔNΔ1Δ2-Ub-prg^-UAF1^ΔSLD^) was concentrated to 4-7 mg/mL and drops of 3-4.5 μL were setup using a reservoir of 8-13% w/v PEG3350, 0.1 M citric acid/Bis-Tris propane pH 4.1 and a ratio of 2 volumes protein to 1 volume reservoir. For cryo-protection, crystals were transferred into reservoir solution supplemented with 25% glycerol and incubated for 5 min up to 12 hours, followed by vitrification in liquid nitrogen.

X-ray diffraction data were collected on a PILATUS 6M-F detector at Diamond Light Source, Beamline I04. For USP1^ΔNΔL1ΔL2-Ub-prg^-UAF1^ΔSLD^, indexing and integration were performed using iMosflm^43^, and scaling and merging using AIMLESS^44^. For USP1^ΔNΔL1ΔL2^-UAF1^ΔSLD^ diffraction was anisotropic and was processed using the STARANISO web server (http://staraniso.globalphasing.org/cgi-bin/staraniso.cgi). For USP1^ΔNΔ1Δ2-Ub-prg^-UAF1^ΔSLD^, molecular replacement was performed using PHASER^45^, with 5CVN chain B (with side chains removed) and 5CVL used as search models for USP1 and UAF1 respectively. One USP1 molecule and two UAF1 molecules were placed. Strong density positive difference density for the Zn atom in the USP1 fingers subdomain was used to validate the molecular replacement solution. Iterative manual modeling building and automated refinement were performed using COOT^46^ and Refmac5^47^, respectively. During automated refinement UAF1 from 51KA (chain B) was used to provide reference restraints. After several rounds of refinement, another USP1 molecule and two ubiquitin molecules were placed using rigid body fitting. For USP1^ΔNΔ1Δ2^-UAF1^ΔSLD^, molecular replacement was performed using USP1^ΔNΔ1Δ2-Ub-prg^-UAF1^ΔSLD^ chains A and B, and refined as above. Final refinement steps were performed in phenix^48^. Crystallography data and model statistics are reported in Supplementary Table S1.

### SAXS data collection and processing

Size-exclusion chromatography (SEC) coupled to SAXS measurements were performed at the Diamond Light Source, Beamline B21 via the mail-in service. USP1^ΔNΔ1Δ2-Ub-prg^-UAF1^ΔSLD^ at ∼10 mg/mL was thawed and injected at 0.075 mL/min onto a Superose 6 Increase 3.2/300 column equilibrated with 20 mM Tris pH 8.0, 200 mM NaCl, 5% glycerol, 5 mM DTT. SAXS data were collected on an Eiger 4M detector. Buffer subtraction and averaging were performed using chromixs^49^. Buffer subtracted, averaged scattering was analyzed using ATSAS^49^ and RAW^50^. Crysol was used to fit PDB models to the SAXS data.

### Gel filtration analysis

To prepare the USP1^C90S^-UAF1-FANCI-FANCD2^Ub^ complex the four individually purified subunits were thawed on ice and mixed 1.2:1.2:1:1. The assembled complex was diluted to ∼5 mg/mL and a total volume of 250 μL with 20 mM Tris pH 8.0, 25 mM NaCl, 5% glycerol, 3 mM MgCl2, 1 mM DTT to yield a final NaCl concentration of approximately 200 mM. The sample was then injected at 0.5 mL/min onto a Superose 6 Increase GL 10/300 column equilibrated in EM buffer (20 mM Tris pH 8.0, 150 mM NaCl, 2 mM DTT).

### Cryo-EM sample preparation and data collection

The USP1^C90S^-UAF1-FANCI-FANCD2^Ub^ complex was prepared by mixing the four individually purified subunits 1:1:1:1. The complex was exchanged into EM buffer using a Bio-Spin P-30 column (Bio-Rad). The concentration of complex was estimated from absorbance at 280 nm and 1 equivalent of dsDNA (61 base pairs; TGATCAGAGGTCATTTGAATTCATGGCTTCGAGCTTCATGTAGAGTCGACGGTGCTGGGAT; IDT) per protein complex was added. Immediately prior to preparing grids the sample was pre-equilibrated to room temperature for 5 min. Quantifoil 1.2/1.3 300 mesh grids were glow discharged twice and 3.6 μL of at 8.8 μM complex was applied. The grids were blotted for 2.5 s and vitrified in liquid ethane using a Vitrobot operating at ∼95% humidity at 15°C.

### Cryo-EM data collection and processing

A CRYO ARM 300 (JEOL) equipped with a DE64 detector was used to collect a total of 4912 movies using beam-Image shift at the Scottish Centre for Macromolecular Imaging (SCMI). All movies were collected in counting mode with a calibrated pixel size of 1.015 Å using SerialEM^51^. Movies were as 59 frames for a total dose of ∼65 e^-^/Å^2^ over 14.9 s. A subset of the movies were additionally gain corrected using relion_estimate_gain^52^. Subsequent processing was performed in Cryosparc v2.13.2^53^ (fig. SB-D). Patch motion correction, patch CTF estimation, and manual curation was performed resulting in 4593 dose-weighted, motion corrected images. Blob picking was performed with minimum and maximum particle diameters of 100 Å and 220 Å respectively, using an elliptical blob. 2D classification was used to generate classes for template picking and select particles for *ab initio* reconstruction with 2 classes, from which a single class with clear protein features was obtained. Template picking was performed on the entire dataset and 1,485,510 boxes were extracted. To remove non-particles, multiple rounds of heterogenous refinement were performed with one good starting model and low pass filtered to 20 Å and 2 or 3 “junk” starting models not representing the protein complex of interest. This cleaning resulted in 391,552 particles for which homogeneous refinement, followed by non-uniform, local refinement using the dynamic mask generated from homogenous refinement, were performed.

The local resolution was performing using FSC threshold of 0.5 and Adaptive Window Factor of 8, which was then used for local filtering. DeepEMhancer (version 0.13)^36^ via COSMIC2^54^ was used to post-process the half-maps using the highRes learning model to aid interpretation.

### Model building

Initially rigid body fitting was performed with USP1 and ubiquitin from the ubiquitin-free structure (chains A and C), UAF1 from 5K1A (chain B), and FANCI and FANCD2 from 6VAF (chains A and B) using UCSF Chimera^55^. The N-terminal region of FANCD2 (residues 45 to 311) was fitted as a rigid body into the locally filtered map^46^. Manual model editing was performed using COOT and incorporating torsion, planar peptide, trans peptide, and Ramachandran restraints. Where appropriate secondary structure restraints were also included. Automated refinement against a globally sharpened map, with the B-factor estimated from the Guinier plot, was performed using phenix real-space refinement^56^. A non-bonded weight of 250 was used to reduce steric clashes. Bond and angle restraints for the USP1 Zinc finger and the K561-G76 isopeptide bond were incorporated. UAF1 from 51KA (chain B) was used to provide reference restraints. The dsDNA was modelled using rigid body fitting of chains S and T from 6VAE. Cryo-EM data and model statistics are reported in Supplementary Table S2.

Superposition of atomic models was performed using “MatchMaker” in UCSF Chimera with the default settings unless otherwise specified. Rotations were estimated using the “measure rotation” command in UCSF Chimera. Buried surface area calculations were performed using PISA^57^. Figures were produced with UCSF Chimera^55^ or ChimeraX^58^.

### Deubiquitination assays

Deubiquitination reactions were performed by preparing a 2x substrate mix and a 2x enzyme mix, and mixing these 1:1 to initiate the reaction. Both mixes were setup on ice, and then incubated at room temperature 30 min prior to reaction initiation and during the reaction. 5 μL aliquots of the reaction were terminated at the indicated timepoints by addition of 5 μL 2x NuPAGE LDS buffer (Thermo Fisher) supplemented with 200 mM DTT.

The 2x substrate mixes were prepared by diluting stocks (≥30 μM) of FANCD2_Ub_, His_6_-V5-TEV-FANCI (or matched buffer), and dsDNA (61 base pairs) with DUB buffer (20 mM Tris pH 8.0, 75 mM NaCl, 5% glycerol, 1 mM DTT). The resulting 2x mixes were composed 2 μM FANCD2_Ub_, 2 μM FANCI, 8 μM dsDNA. The 2x enzyme mixes were prepared by diluting concentrated stocks (≥30 μM) of USP1, and UAF1 (full-length or β-propeller domain) with DUB buffer. The resulting 2x mixes were composed of 200 nM USP1 and 200 nM UAF1 (full-length) or 400 nM USP1 and 400 nM UAF1 (β-propeller domain). SDS-PAGE, Coomassie staining, and western blotting was performed as described previously ^18^. For western blotting, 1:1000 dilution Anti-FANCD2 (ab108928) and 1:8000 dilution Anti-V5 (ab27671; 1 in 8000 dilution) were used for detection of FANCD2 and FANCI, respectively. After labelling secondary antibodies at 1:10,000 dilution, bands were visualized on an Odyssey CLX Infrared Imaging System (Li-COR). Each band was quantified using Image Studio with horizontal background subtraction (Li-COR). Replicate assays were performed from different thawed protein aliquots, each from the same protein preparation.

## Data availability

The atomic coordinates and structure factors of ubiquitin-free and ubiquitin-bound USP1-UAF1 are being deposited to the PDB. The atomic coordinates and cryo-EM maps, including locally filtered and sharpened and DeepEMhancer maps, are being deposited to the PDB and EMDB.

## Acknowledgements

We thank past and current members of the Walden laboratory for experimental suggestions, comments on the manuscript and their support. All constructs are available on request from the MRC Protein Phosphorylation and Ubiquitylation Unit reagents Web page (http://mrcppureagents.dundee.ac.uk) or from the corresponding author. We acknowledge Diamond Light Source for time on beamline I04 (proposal mx14980) and B21 (proposal mx19844) and thank Dr. Nikul Khunti for collecting SAXS data. We acknowledge the Scottish Centre for Macromolecular Imaging (SCMI) for access to cryo-EM instrumentation, funded by the MRC (MC_PC_17135) and SFC (H17007), and thank Dr. Mairi Clarke and Dr. James Streetley for screening and collection of cryo-EM data. This work was supported by the EMBO Young Investigator Programme to H.W.; the European Research Council (ERC-2015-CoG-681582 ICLUb consolidator grant to H.W.

## Author contributions

M.L.R., C.A., V.K.C. and H.W. conceived this work; C.A., M.L.R. and V.K.C. purified proteins; C.A. and R.T. generated various expression vectors and performed mutagensis; C.A. performed crystallography; M.L.R. performed cryo-EM data processing; M.L.R. and C.A. performed model building and refinement; M.L.R. performed assays; M.L.R. and C.A. wrote the manuscript with contributions from all other authors; H.W. secured funding and supervised the project.

