## Supplementary Figures and Tables for "Structural basis of FANCD2 deubiquitination by USP1-UAF1"

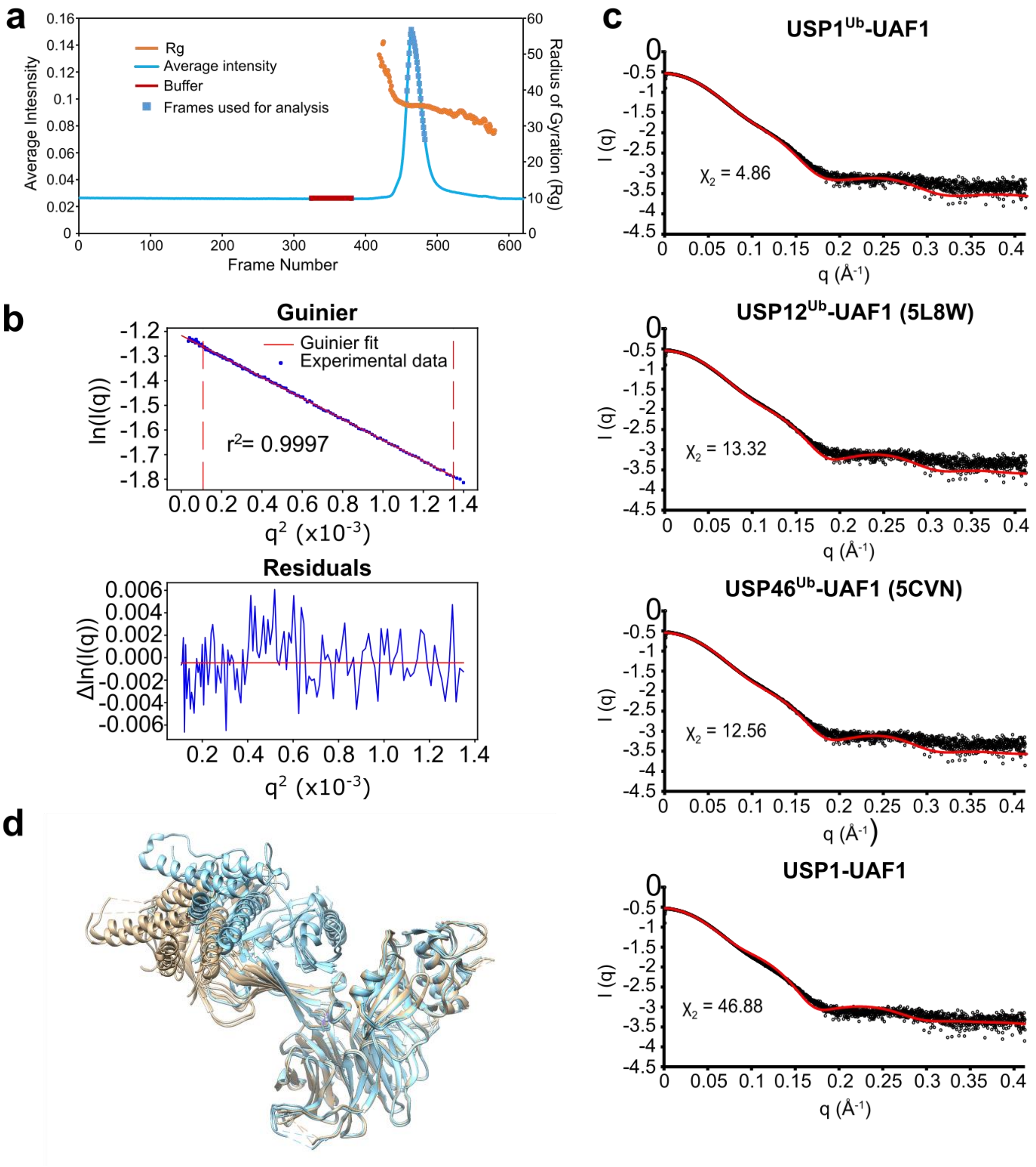


**Supplementary Figure S1.** Structural characterization of USP1-UAF1. (**a**) SEC-SAXS trace for the crystallized ubiquitin-bound USP1-UAF1 construct. (**b**) Guinier plot of buffer subtracted, averaged SAXS measurements. (**c**) Fit of different USP-UAF1 crystal structures to SAXS measurements. The better resolved chains A and B of the ubiquitin-free (USP1-UAF1) and chains A, B, and C ($USP1^{Ub}$-UAF1) of the ubiquitin-bound structure were used for fitting. (**d**) The two USP1-UAF1 complexes in the asymmetric unit of the ubiquitin-free (brown) and ubiquitin-bound (blue) crystal structures aligned by UAF1. The better resolved chains A and B of the ubiquitin-free (USP1-UAF1) and chains A, B, and C USP1$^{Ub}$-UAF1) of the ubiquitin-bound structure were used for analysis unless otherwise stated.

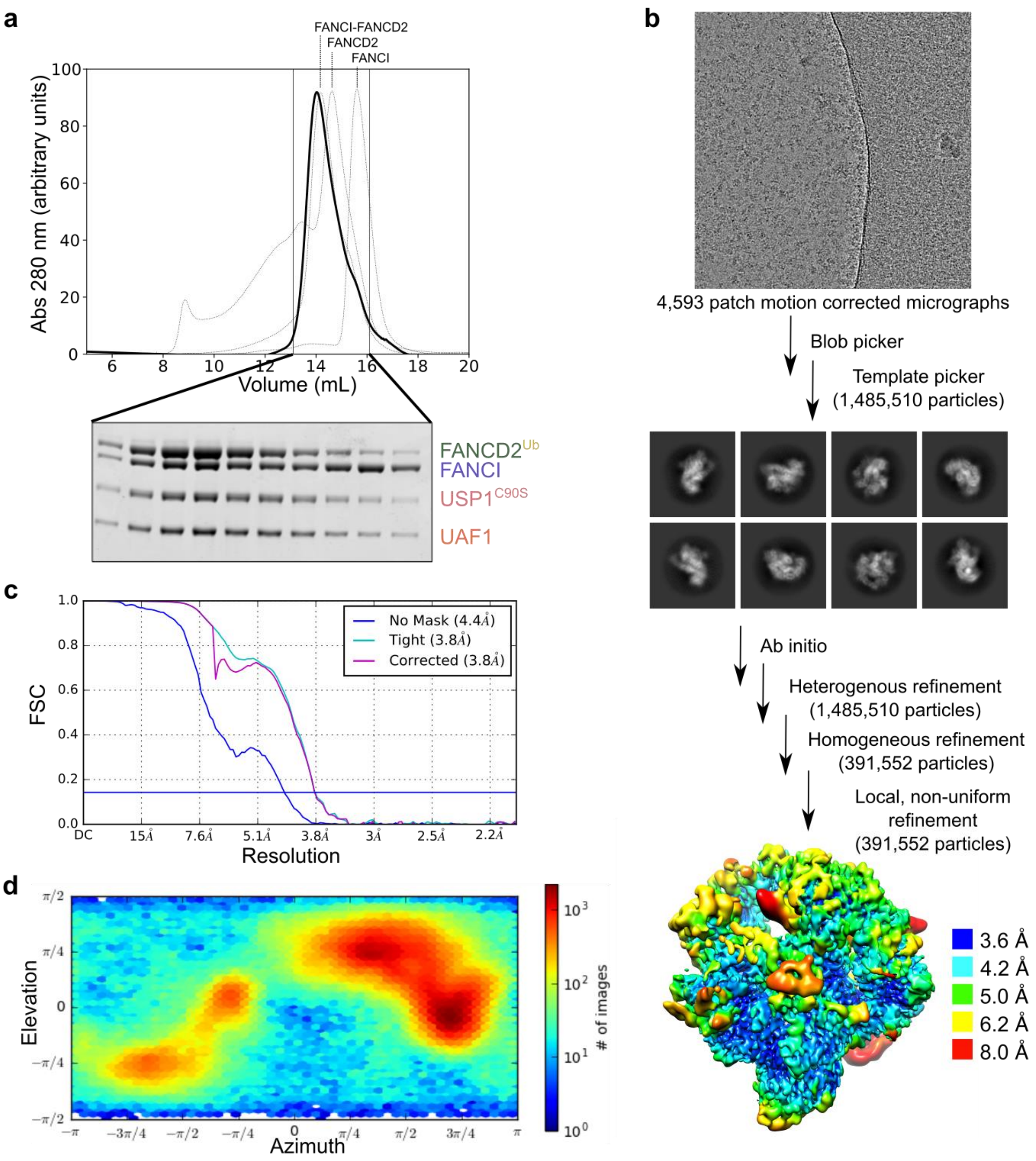


**Supplementary Figure S2.** Cryo-EM data processing. (**a**) Gel filtration profile of the assembled USP1$^{C90S}$-UAF1-FANCI-FANCD2$^{Ub}$ complex and associated SDS-PAGE and Coomassie staining. (**b**) Flow diagram for analysis of the cryo-EM data by single particle analysis (see also Methods). The locally filtered map is shown. (**c**) Unmasked, masked, and corrected Fourier Shell Correlation curves. (**d**) Viewing direction distribution.

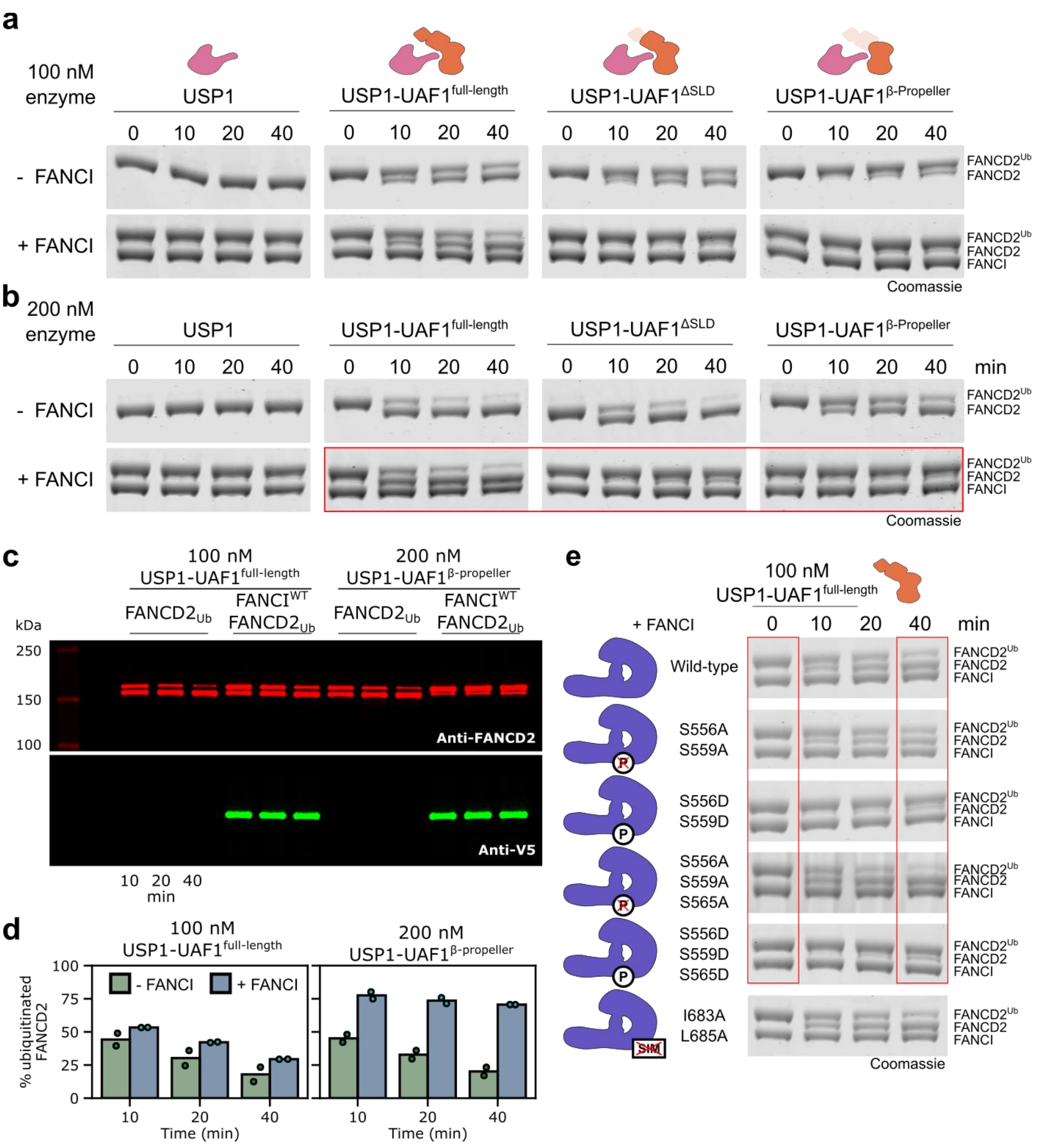


**Supplementary Figure S3.** Deubiquitination assays of FANCD2 by USP1-UAF1. (**a**) Deubiquitination time-courses for full-length USP1 alone and with the addition of various UAF1 truncations, at 100 nM USP1, 100 nM UAF1 as assessed by SDS-PAGE and Coomassie staining. (**b**) Deubiquitination time-courses for USP1 alone and with the addition of various UAF1 truncations, at 200 nM USP1, 200 nM UAF1 as assessed by SDS-PAGE and Coomassie staining. (**c**) Deubiquitination time-courses as assessed by Western blot. (**d**) Quantification of Western blots. Mean values are represents as bars with the individual replicates shown as points. (**e**) Deubiquitination time-courses for full-length USP1 and full-length UAF1 with ubiquitinated FANCD2 and various FANCI mutants as assessed by SDS-PAGE and Coomassie staining. For all assays 1 µM ubiquitinated FANCD2 was used, and 1 µM FANCI where included. All assays were in the presence of 4 µM 61 base pair dsDNA, and performed at least twice (two technical replicates). Red boxes indicate results displayed in Figure 3.

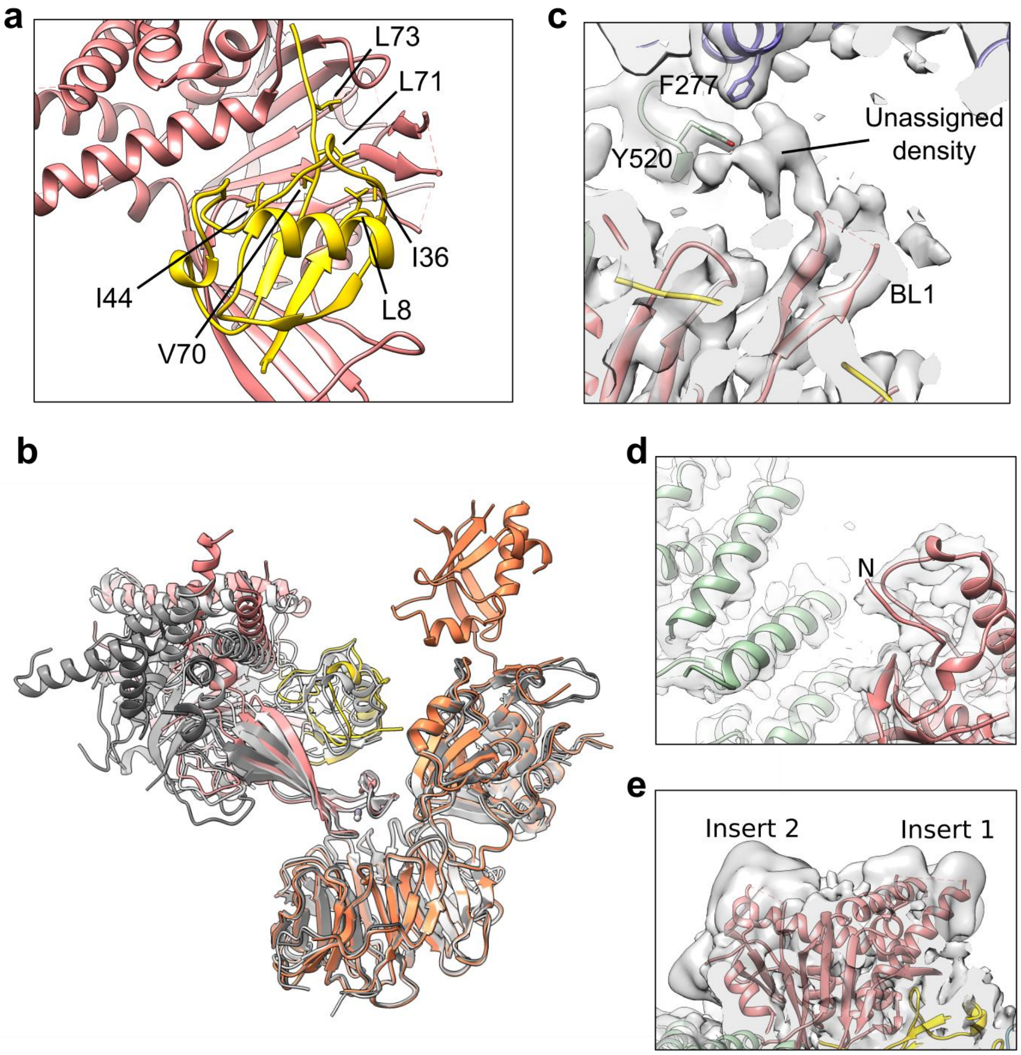


**Supplementary Figure S4.** The structure of USP1-UAF1 when bound to FANCI-FANCD2$^{Ub}$. (**a**) The USP1-ubiquitin interface. Residues of the hydrophobic patch of ubiquitin are highlighted. (**b**) Alignment of

USP1-UAF1 structures by the UAF1 subunit. The ubiquitin-free (dark gray) and ubiquitin-bound (light gray) crystal structures, and the FANCI-FANCD2$^{Ub}$-bound cryo-EM structure (colored) are shown. (**c**) An unassigned blob of density is present adjacent to FANCI, FANCD2 and the BL1 of USP1. The locally filtered map is shown at a threshold of 0.07. (**d**) The N-terminus of USP1 is not well resolved. The DeepEMhancer map is shown at a threshold of 0.1. (**e**) Inserts 1 and 2 of USP1 are not well resolved. The locally filtered map is shown at a threshold of 0.02.

**Supplementary Table S1.** Crystallography data collection and model refinement statistics[a]

| | Ubiquitin-free USP1-UAF1 | Ubiquitin-bound USP1-UAF1 |
|---|---|---|
| **Data collection and processing** | | |
| Beamline | DLS I04 | DLS I04 |
| Wavelength (Å) | 0.9795 | 0.9795 |
| Unit cell | a=b=119.57 Å, c=195.46 Å | a=b=134.24 Å, c=274.67 Å |
| | α=β=γ=90° | α=β=90°, γ=120° |
| Space group | $P4_1$ | $P6_5$ |
| Resolution range (Å) | 84.549 – 3.602 (3.754 – 3.602) | 91.56 – 3.20 (3.31 – 3.20) |
| Ellipsoidal resolution (Å) (direction) | 4.312 (0.902 a* + 0.433 b*) | N.A. |
| | 4.381 (0.879 a* + 0.476 b*) | |
| | 3.601 (0.246 a* + 0.469 b* + 0.848 c*) | |
| Unique reflections | 25470 (1264) | 46123 (4501) |
| Multiplicity | 5.7 (5.4) | 10.5 (10.1) |
| Completeness (%) | | |
| Spherical | 80.3 (34.2) | 100 (100) |
| Ellipsoidal | 95.0 (100) | N.A. |
| Mean I/sig(I) | 6.4 (1.7) | 11.6 (1.7) |
| Wilson B-factor (Å$^2$) | 102.7 | 62.8 |
| $R_{meas}$ | 0.258 (1.698) | 0.193 (1.657) |
| $R_{pim}$ | 0.108 (0.727) | 0.059 (0.519) |

| | | |
|---|---|---|
| $CC_{1/2}$ | 0.992 (0.565) | 0.997 (0.581) |
| **Refinement** | | |
| $R_{work}$ / $R_{free}$ | 0.235 / 0.266 | 0.204 / 0.234 |
| Number molecules in asymmetric unit | | |
| USP1 | 2 | 2 |
| UAF1 | 2 | 2 |
| Ubiquitin | 0 | 2 |
| Disorder model | One TLS group per protein molecule + residual isotropic B-factors | One TLS group per protein molecule + residual isotropic B-factors |
| Bond length rmsd (Å) | 0.003 | 0.004 |
| Bond angle rmsd (°) | 0.808 | 0.909 |
| All-atom clashscore | 7.3 | 3.9 |
| Ramachandran plot | | |
| Outliers (%) | 0.51 | 0.31 |
| Favored (%) | 93.79 | 95.91 |
| Rotamer outliers (%) | 1.39 | 0.07 |
| PDB ID | 7AY0 | 7AY2 |

[a]Values in parentheses correspond to the highest resolution shell, apart from Ellipsoidal resolution

**Supplementary Table S2.** Cryo-EM data collection and model refinement statistics

| **Data collection and processing** | |
|---|---|
| Microscope | Cryoarm |
| Detector | DE64 |
| Magnification | 120,000x |
| Voltage (kV) | 300 |
| Electron Dose ($e^-$/Å$^2$) | 65 |
| CTF Estimated Defocus range (µm) | 0.22 – 3.7 |

| | |
|---|---|
| Pixel Size (Å) | 1.015 |
| Symmetry imposed | C1 |
| Consensus map resolution (Å) | 3.8 |
| FSC threshold | 0.143 |
| Map resolution range (Å)[a] | 3.6-10 |
| FSC threshold | 0.5 |
| EMDB ID | EMD-11934 |
| **Refinement** | |
| Initial models used | 6VAF, 5K1A, ubiquitin-bound USP1 |
| Map sharpening B-factor ($Å^2$) | 94.6 |
| Correlation coefficient (mask)[b] | 0.83 |
| Bond length rmsd (Å) | 0.006 |
| Bond angle rmsd (°) | 0.973 |
| All-atom clashscore | 9.1 |
| Ramachandran plot | |
| Outliers (%) | 0.04 |
| Favored (%) | 93.61 |
| Rotamer outliers (%) | 0.07 |
| PDB ID | 7AY1 |

[a]1% and 99% quantiles

[b]Calculated in phenix

**Supplementary Movie S1.** Morph between FANCI-FANCD2$^{Ub}$ on its own (6VAF) and bound to USP1-UAF1. FANCD2 – green; FANCI – violet; ubiquitin – yellow; USP1-UAF1 – transparent.

**Supplementary Movie S2.** 3D variability of the cryo-EM dataset of USP1-UAF1-FANCI-FANCD2$^{Ub}$. First eigenvector of variability in the dataset. Viewed from above with respect to Figure 2, with FANCD2 on the left hand side.

**Supplementary Movie S3.** 3D variability of the cryo-EM dataset of USP1-UAF1-FANCI-FANCD2$^{Ub}$. Second eigenvector of variability in the dataset. Viewed from above with respect to Figure 2, with FANCD2 on the left hand side.
